# SULFATED POLYSACCHARIDE FROM BROWN ALGAE *DICTYOTA CARIBAEA* STIMULATES MACROPHAGES TO AN ANTITUMORAL PHENOTYPE

**DOI:** 10.1101/2023.06.15.545132

**Authors:** Alexia Nathália Brígido Assef, Felipe Barros Teles, Leonardo Paes Cinelli, Diego Veras Wilke

## Abstract

Fucoidans are sulfated polysaccharides capable of exerting biological activities such as antitumor and immunomodulatory effects. Previous studies demonstrated the antitumor activity of non-cytotoxic fucoidan from the seaweed *Dictyota caribaea* (Dc-SP) *in vivo*. Macrophages (Mφ) are innate immune cells capable of promoting or inhibiting tumor growth depending on the stimulus. This study aimed to evaluate the immunostimulant activity of Dc-SP on murine Mφ (RAW264.7) *in vitro*. Dc-SP was assessed for its ability to modify cell viability and stimulate the production of antitumor markers on RAW264.7 cells. Dc-SP induced an increase (p<0.05) in the production of NO and cytokines TNF-α, IL-1β, and IL-10. The exposure of Mφ to Dc-SP also increased (p<0.05) the expression of M1 macrophage markers such as iNOS, CD86, and MHC II. The antiproliferative activity of RAW264.7 cells stimulated with DC-SP on melanoma cells (B16-F10) *in vitro*. Dc-SP did not exhibit direct cytotoxicity on B16-F10, however, the conditioned medium (CM) of RAW264.7 previously stimulated with Dc-SP (CM-Dc-SP) showed antiproliferative activity on tumor cells. B16-F10 incubated with CM-Dc-SP showed a cytostatic profile, tumor cells did not alter membrane integrity, however, suffered morphological changes such as cell shrinkage and high granularity. In conclusion, Dc-SP stimulated Mφ to an antitumor phenotype.

## 1. Introduction

Mφ are innate immune cells involved in a myriad of biological processes. Mφ play different roles in the development of cancer and may present a phenotype with antitumor activity in the early stages of progression and even another phenotype capable of promoting tumors in established cancer [1]. Notably, Mφ are highly plastic cells and, depending on stimulation in the tumor microenvironment (TME), may undergo phenotypic changes in their function. Often, high infiltration of Mφ is strongly associated with a worse prognosis for the patient or even a tumor progression in many types of solid tumors, including melanoma [2]. In addition, Mφ can present antigens, secrete several important molecules, and can regulate the tumor microenvironment and other physiological functions [3]. Depending on the stimulus, these Mφ can be polarized to a classic activated Mφ (M1) or an alternative activated Mφ (M2) phenotype with different marker expressions involved [4]. Tumor-associated Mφ M2 type (TAMs) are responsible for tissue repair, tumor angiogenesis, and wound healing, and contribute to tumor growth [5,6]. Therefore, it is beneficial to have higher levels of M1 Mφ, which have an anti-tumoral phenotype.

Natural products, including polysaccharides, have been studied for their diverse biological effects such as antitumor and immunomodulatory activities [7]. Thus, they are considered important biological response modulators (BRM) capable of activating the host’s innate and acquired immune systems [8]. Fucoidans are natural polysaccharides containing L-fucose and sulfate ester groups. Also called homofucans, they are constituents of brown seaweed and some marine invertebrates (such as sea urchins and sea cucumbers) [9]. Fucoidans show antitumor properties and immunomodulatory effects with low toxicity, anti-inflammatory, and other pharmacological properties [10-12]. In our previous studies, we demonstrated that fucoidan from the brown algae Dc-SP exhibited an inhibitory effect *in vivo* on the sarcoma-180 tumor. However, it did not induce cytotoxicity on tumor cells *in vitro*. Additionally, Dc-SP did not show toxic effects on mice. Nevertheless, spleen weight increased along with an activated morphology, suggesting the observed antitumor effect may be related to an immunomodulatory effect of Dc-SP [13]. *In vitro* studies are suitable to accelerate drug discovery in several aspects highlighting the potential use of substances [14,15]. They include low cost and amount of substances, reproducibility, and avoidance of ethical issues. In this work, we evaluated the MΦ activation *in vitro* to an antitumor phenotype on RAW264.7 cells exposed to Dc-SP.

## 2. Materials and methods

### 2.1. Reagents

Dulbecco’s modified Eagle medium (DMEM), fetal bovine serum (FBS), antibiotics (penicillin and streptomycin), and trypsin-EDTA were purchased from Gibco BRL Co. (Grand Island, NY, USA). Lipopolysaccharide from *Escherichia coli* (LPS), acetic acid, triton X-100, sulfanilamide, and NED reagents was obtained from Sigma Aldrich (St. Louis, MO, USA). Sodium nitrite (NaNO_2_) was purchased from Dinâmica (Diadema, São Paulo, Brazil). TNF-α, IL-1β, and IL-10 ELISA kit was purchased from R&D Systems (Minneapolis, MN, USA). Surface markers anti-CD86, anti-MHCII, and intracellular marker anti-iNOS were purchased from Thermo Fisher (Waltham, MA, USA).

### 2.2. Isolation of sulfated polysaccharide from *Dictyota caribaea (Dc-SP)*

The marine algae, *Dictyota caribaea*, was collected in March 2011 at Vermelha beach (23°11’46.0” S and 44°38’38.0” W), Paraty, Rio de Janeiro, Brazil, separated of epiphytes and from other species, washed with distilled water, air-dried, powdered and stored at -20°C. Voucher specimens (RFA 38779) were deposited at the Herbarium of the Rio de Janeiro Federal University, Brazil (Simas et al, 2014). The use of *D. caribaea* was registered on the National System of Management of Genetic Heritage and Associated Traditional Knowledge (SisGen) under the number #AC2D331. Dc-SP was obtained by proteolytic digestion using papain incubated at 60°C for 12 h and sulfated polysaccharide was precipitated in the presence of ethanol (at final concentrations of 9%) to obtain Dc-SP. The peak obtained at DEAE-cellulose (Dc-SP) was applied to a High Q-HPLC at the same conditions of equilibration, elution, collection, and fraction checked as previously described [13].

### 2.3. Cell lines, cell culture procedures, and treatments

RAW264.7, a murine MΦ cell line, and B16-F10, a murine metastatic melanoma cell line, were purchased from the Banco de Células do Rio de Janeiro (Rio de Janeiro, Brazil). They were grown with DMEM medium, and supplemented with 10% fetal bovine serum, 2 mM glutamine, 100 μg/mL streptomycin, and 100 U/mL penicillin and incubated at 37 °C with a 5% CO_2_ atmosphere.

The Mφ activation assays were conducted with RAW264.7 cells plated at 1,5 x 10^5^ cells/mL and incubated for 24 h with saline (as negative control), *Escherichia coli* lipopolysaccharide at 100 ng/mL (LPS) as a positive control, or Dc-SP at 10, 100, and 250 µg/mL. The antiproliferative assays were carried out with B16-F10 cells plated at 4 x 10^4^ cells/mL and incubated for 48h with conditioned medium obtained from saline-treated Mφ (CM-Saline), LPS-treated Mφ (CM-LPS), Dc-SP-treated Mφ (CM-Dc-SP) or Doxorubicin at 0.6 µM (DOX) as a gold standard antiproliferative positive control.

### 2.4. Griess assay

Mφ activation was initially determined by evaluating the NO production. Griess assay was performed to measure nitrites accumulation the product of NO metabolism [16]. RAW264.7 cells were incubated for 24 h with saline, LPS 100 ng/mL, or Dc-SP (at all concentrations described above). The nitric oxide (NO) levels formed in the supernatant of Mφ RAW264.7 were indirectly measured by the Griess test.

### 2.5. Cytokines expression

The levels of cytokines IL-10, TNF-α, and IL-1β present in the CM samples were quantified by ELISA using antibodies obtained from R&D Systems according to the manufacturer’s instructions.

### 2.6. Flow cytometry analysis

We used FITC-conjugated anti-CD86 (11-0862-82) and PE-conjugated anti-MHCII (12-5320-82) antibodies from Thermo Fisher (Waltham, MA, USA) to stain the Mφ activation markers CD86 and MHCII, respectively. Cells were incubated with the antibodies for 30 minutes at 4°C in FACS buffer (PBS supplemented with 4% FBS) and then washed with FACS buffer again. The cells were fixed with paraformaldehyde 1% for 5 minutes and permeabilized with Triton 0.1% for 5 minutes for iNOS labeling. Then, cells were incubated with Alexa Fluor 488-conjugated anti-NOS2 (53-5920-82) for 30 minutes at 4 ºC in FACS buffer and washed with FACS buffer again. We used flow cytometry (model FACSVerse, BD Biosciences, San Jose, USA) for data acquisition and FlowJo software (San Jose, CA, USA) for the analysis. Ten thousand events were acquired excluding debris and doublets for each tube. Median fluorescence intensity values (MFI) of iNOS, MHCII, and CD86 staining were normalized using the mean MFI values of the negative control. Additionally, the percentage of double-labeled cells was used to show the population of cells expressing MHC II and CD86 simultaneously.

### 2.7. Evaluation of antiproliferative effect in vitro

The antiproliferative effect of samples was performed by the sulforhodamine B (SRB) assay [17]. SRB assay is used for the determination of cell density, based on the measurement of total cellular protein content, and does not depend on cellular metabolism.

B16-F10 cells were then incubated for 48h with CM-Saline, CM-LPS, CM-Dc-SP, or DOX. Therefore, we added 100 µL of CM to an equal volume of the B16-F10 supernatant (1:1 v/v), resulting in a dilution factor = 2. Absorbances were read on a plate spectrophotometer (Fisher Scientific, model Multiskan FC) at 564 nm or 570 nm. Using the same treatments above, Flow cytometry experiments were performed to evaluate changes in cell characteristics. Also, cell count, membrane integrity, and morphological changes were assessed. Flow cytometry analysis was performed using appropriate gating strategies for the desired cell population, with debris and aggregates excluded from the analysis. 10 thousand events were acquired in the flow cytometer for evaluation of cell count, morphology, and cell viability. The results were analyzed using FlowJo software.

### 2.8. Statistical analyses

The data are expressed as mean ±SD. The surface staining for iNOS quantification and the morphological parameters assay were performed in two independent experiments. All other experiments were performed in three independent experiments. The differences between experimental groups were compared by One-way Analysis of Variance (ANOVA) followed by Dunnet’s post-test to evaluate the tumor and organs weight (p<0.05) and Newman-Keuls’ post-test for evaluating the biochemical and hematological parameters (P<0.05) using the GraphPad PRISM program 5 (Intuitive Software for Science, San Diego, CA, USA).

## 3. Results and Discussion

### 3.1. Dc-SP polarizes macrophages towards to M1 phenotype

#### 3.1.1 Nitric Oxide and Cytokines Release

Dc-SP stimulated the production of NO levels (Fig 1A) and increases iNOS (Fig 2A-B) expression in RAW264.7 cells. In addition, Dc-SP also increased secretion of the cytokines TNF-α, IL-1β, and IL-10 (Fig 1B-D), when compared to the negative control group. Nitric oxide is a highly volatile free radical that participates in various physiological and pathological processes in the body, such as the regulation of blood pressure, inhibition of leukocyte adhesion to the endothelium, and induction of apoptosis [18,19]. Both the presence of pathogenic microorganisms and certain substances can activate Mφ, resulting in the secretion of cytokines that play a crucial role in the host immune response [20]. As described in the literature, sulfated polysaccharides have immunomodulatory potential in cytokine production [21-23].

**Fig. 1.**
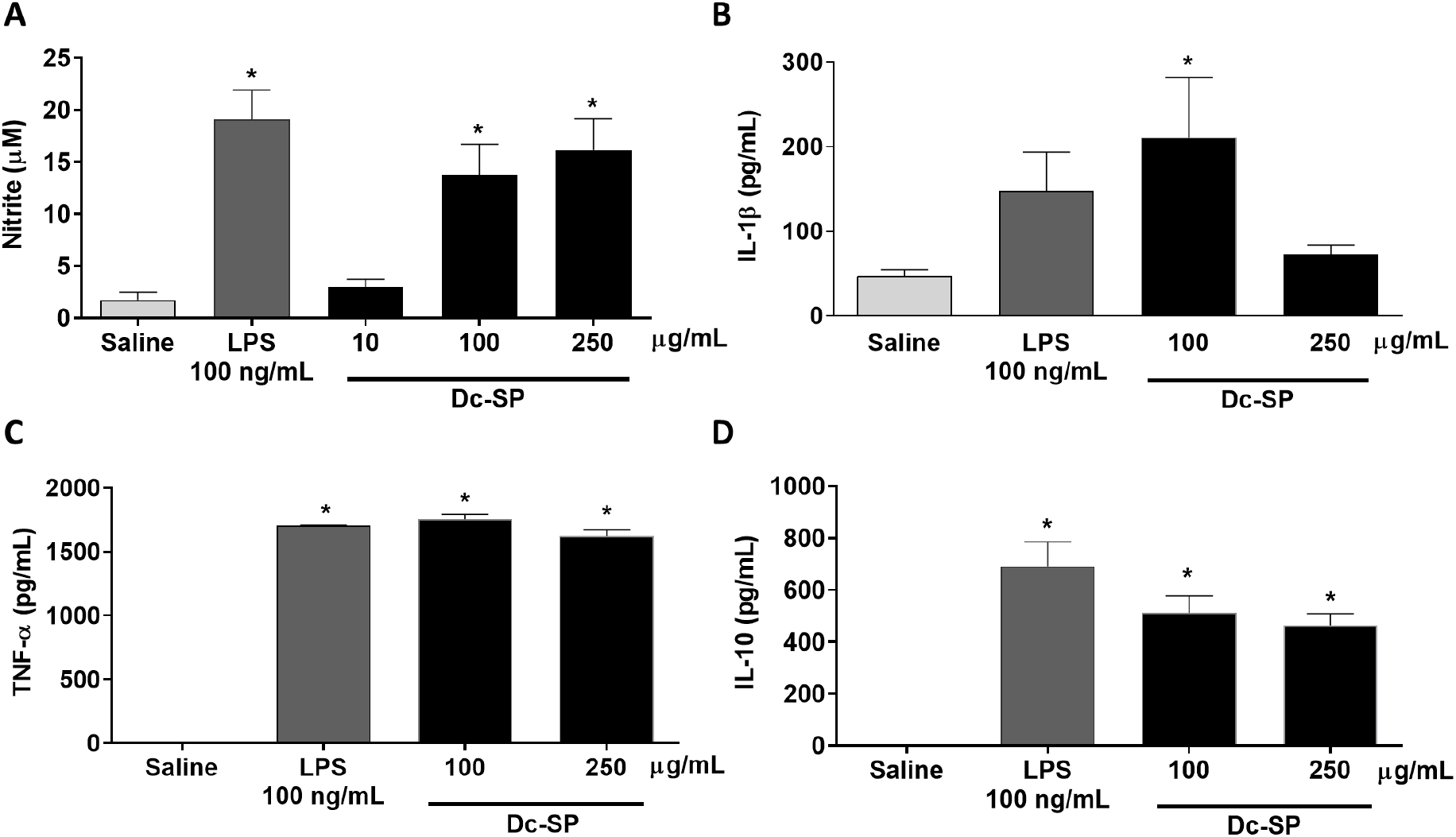
Sulfated polysaccharide from brown algae *Dictyota caribaea* induces an M1 phenotype on macrophages. RAW264.7 cells were exposed to saline as a negative control, *E. coli* lipopolysaccharide at 100 ng/mL (LPS) as a positive control, and increasing concentrations of sulfated polysaccharides from *D. caribaea* (Dc-SP) for 24h. **A**, Nitrite levels produced by Mφ (RAW264.7 cell line) were evaluated by Griess assay, while **B-D** show the levels of IL-β, TNF-α, and IL-10 respectively evaluated by ELISA. Data are presented as mean ±SD. Differences between the negative control versus the other groups were determined by the analysis of variance (ANOVA) followed by Dunnett’s post-test.^*^p<0.05 when compared with negative control. Results are representative of three independent experiments, each one performed in triplicate.

**Fig. 2.**
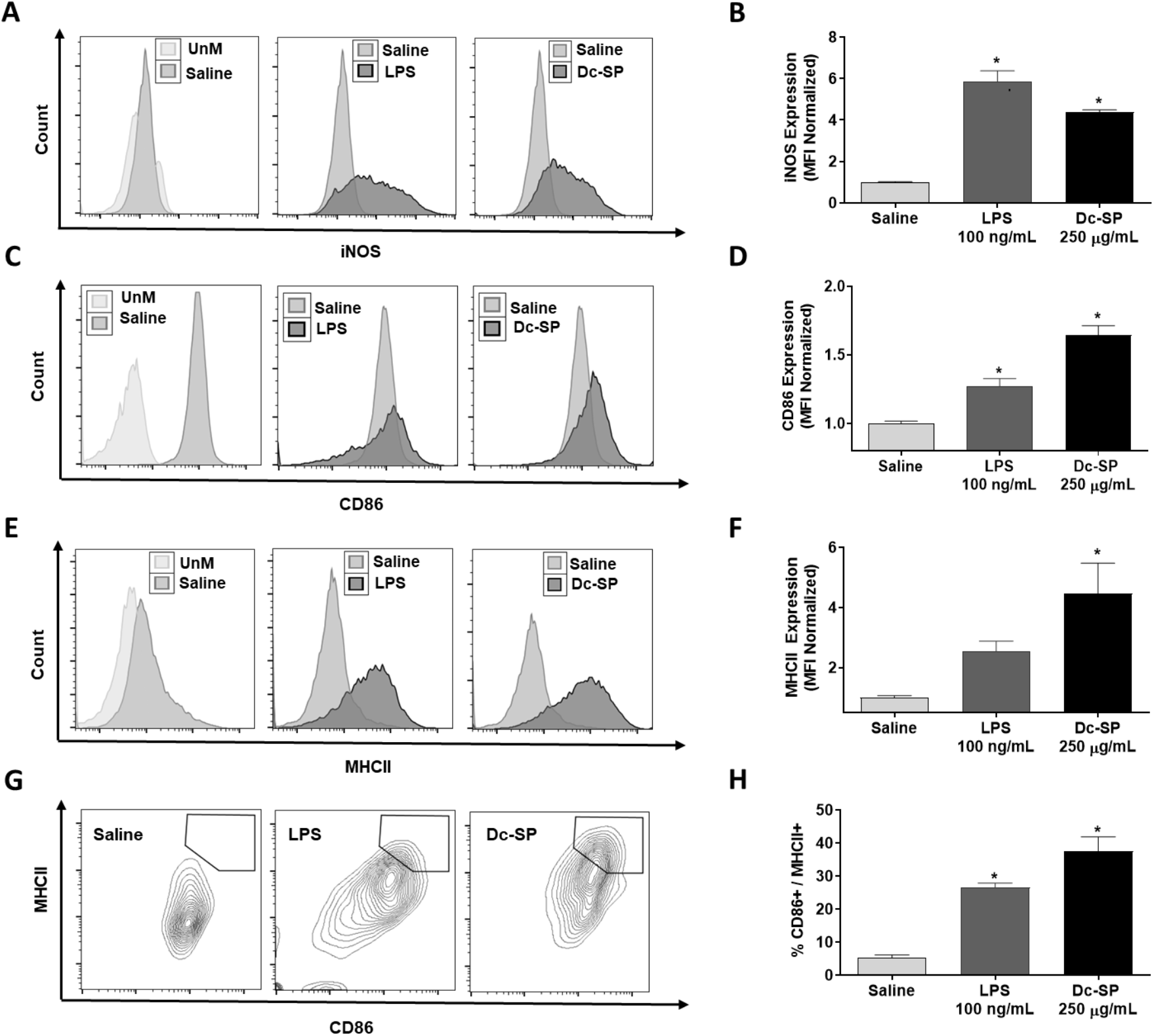
Sulfated polysaccharide from brown algae *Dictyota caribaea* induces expression surface and intracellular markers. Expression of iNOS, CD86, and MHCII was evaluated on RAW264.7 cells by flow cytometry. RAW264.7 cells were incubated for 24 h with saline as a negative control, *E. coli* lipopolysaccharide at 100 ng/mL (LPS) as a positive control, and Dc-SP at 250 µg/mL. Flow cytometry histograms are depicted in **A, C**, and **E**. Median fluorescence intensity (MFI) data are depicted on **B, D**, and **F**, as normalized values. The percentage of CD86^+^/MHCII^+^ cells is depicted in **G** and **H**. The flow cytometry experiments acquired ten thousand events, excluding debris and doublets. Unstained cells (UnM) were used as a reference for basal antibody background. Data on **B, D, F** and **H** are presented as mean ±SD. The mean differences between the negative control versus other groups were compared by analysis of variance (ANOVA) followed by Dunnett’s post-test. Results are representative of three independent experiments, each one performed in triplicate.

Studies have shown that fucans are capable of modulating cytokines and NO, demonstrating the immunomodulatory potential of this polysaccharide [24,25]. Together, these markers indicate that Mφ were polarized towards an M1-like phenotype, which is associated with inflammatory, microbicidal, and tumoricidal activities [26,27]. The immunomodulatory effect of fucoidans on Mφ is associated with the production of pro-inflammatory cytokines, such as TNF-α, IL-6, IL-1β, as well as the production of nitric oxide (NO) and iNOS. [7, 28-30]. Sulfated fucans derived from *Agarum cribrosum* led to a notable elevation in nitric oxide production and the upregulation of iNOS expression and IL-10 because of nuclear factor kappa B (NF-kB) activation in the RAW 264.7 cell culture [31]. Another study shows increased NO production in RAW264.7 in a dose-dependent manner when exposed to fucoidan from brown algae *Nizamuddinia zanardinii*. Moreover, this fucoidan induces an up-regulation of iNOS, TNF-α, IL-6, and IL-1β mRNA [30]. Similarly, fucoidan isolated from *Laminaria angustata* induced Mφ activation to M1-like phenotype since increased NO production and TNF-α and IL-6 synthesis and secretion in RAW264.7 [32]. Fucoidan extracted from *Laminaria japonica* induced the release of NO, IL-6, and TNF-α in RAW264.7. Furthermore, fucoidan from *L. japonica* enhanced the percentage of CD86+ cells, while the percentage of CD206^+^ cells was decreased after incubation with RAW264.7 or mouse bone marrow-derived monocytes (BMDMs) [33].

Tabarsa and cols showed that a fucoidan from *Nizamuddinia zanardinii* upregulated the iNOS expression and NO production and released several pro-inflammatory cytokines such as TNF-α, IL-1β, IL-6, and anti-inflammatory IL-10 [30]. The IL-10 produced inhibited the overexpression of cytokines TNF-α, IL-1β, and IL-6, exhibiting immunological responses [34]. These results suggest that the production of anti-inflammatory cytokines may suppress the overactivation of RAW264.7 Mφ by secreting anti-inflammatory cytokines such as IL-10, thus helping to balance the ratio of pro- and anti-inflammatory cytokines in fucoidan-activated Mφ.

#### 3.1.2 Expression of surface activation markers

Dc-SP increased the expression of the MHC II and the CD86 on the surface of RAW-264.7 cells (Fig. 2C-H). CD86 expression increased by 1.6-fold and about 4.5-fold for MHC-II when compared with the saline group (p<0.05). CD86 is a co-stimulatory molecule considered a marker for M1 Mφ [20,35]. Furthermore, Mφ present antigens to helper T cells through the CD86 and MHC II, regulating innate and adaptive immune responses in the presence of pathogens or activating substances [36]. The up-regulation of MHC II is implicated in antigen presentation and T cell activation, as well as being involved in the interaction between T cells and antigen-presenting cells, such as Mφ, for antitumor activities. [37,38].

A study demonstrated that Polysaccharide-Protein Complex from *Lycium barbarum* L. was able to activate Mφ by inducing the expression of co-stimulatory molecules such as CD80, CD86, and CD40, as well as increasing the production of MHC II and cytokines TNF-α, IL-1β in RAW264.7 Mφ [39]. A polysaccharide from *Tinospora cordifolia* activated Mφ through the classical pathway in a TLR4-MyD88 dependent manner, leading to the production of M1 markers such as nitric oxide (NO), inducible nitric oxide synthase (iNOS), pro-inflammatory cytokines such as TNF-α, IL-β, IL-6, IL-12, IL-10, and IFN-γ, as well as activated Mφ surface markers such as CD86 and MHCII [41]. M1 Mφ activation is advantageous for the host, as cytokines such as IFN-γ, IL-1β, and IL-6 produced by Mφ exhibit an anti-tumoral response. On the other hand, tumor-associated Mφ (TAMs) with an M2 phenotype is often associated with pro-tumoral effects, such as tumor growth, metastasis, and angiogenesis [41,42].

Altogether, our results suggest that Dc-SP is a promising immunomodulatory agent by modulating Mφ to an M1 phenotype. Additionally, Dc-SP exerts *in vivo* antitumor inhibition and may modulate the tumor microenvironment targeting Mφ [13]. Then we further investigate the antiproliferative activity of CM-RAW exposure to Dc-SP on tumor cells, described below.

### 3.2. Antitumor potential of macrophages stimulated with Dc-SP

#### 3.2.1 Conditioned medium obtained from RAW264.7 cells stimulated with Dc-SP inhibits the proliferation of B16-F10 cells

Dc-SP did not show cytotoxicity against the B16-F10 murine melanoma cells (Data not shown). However, the CM samples showed inhibition of tumor growth cells (Fig. 3A).

**Fig. 3.**
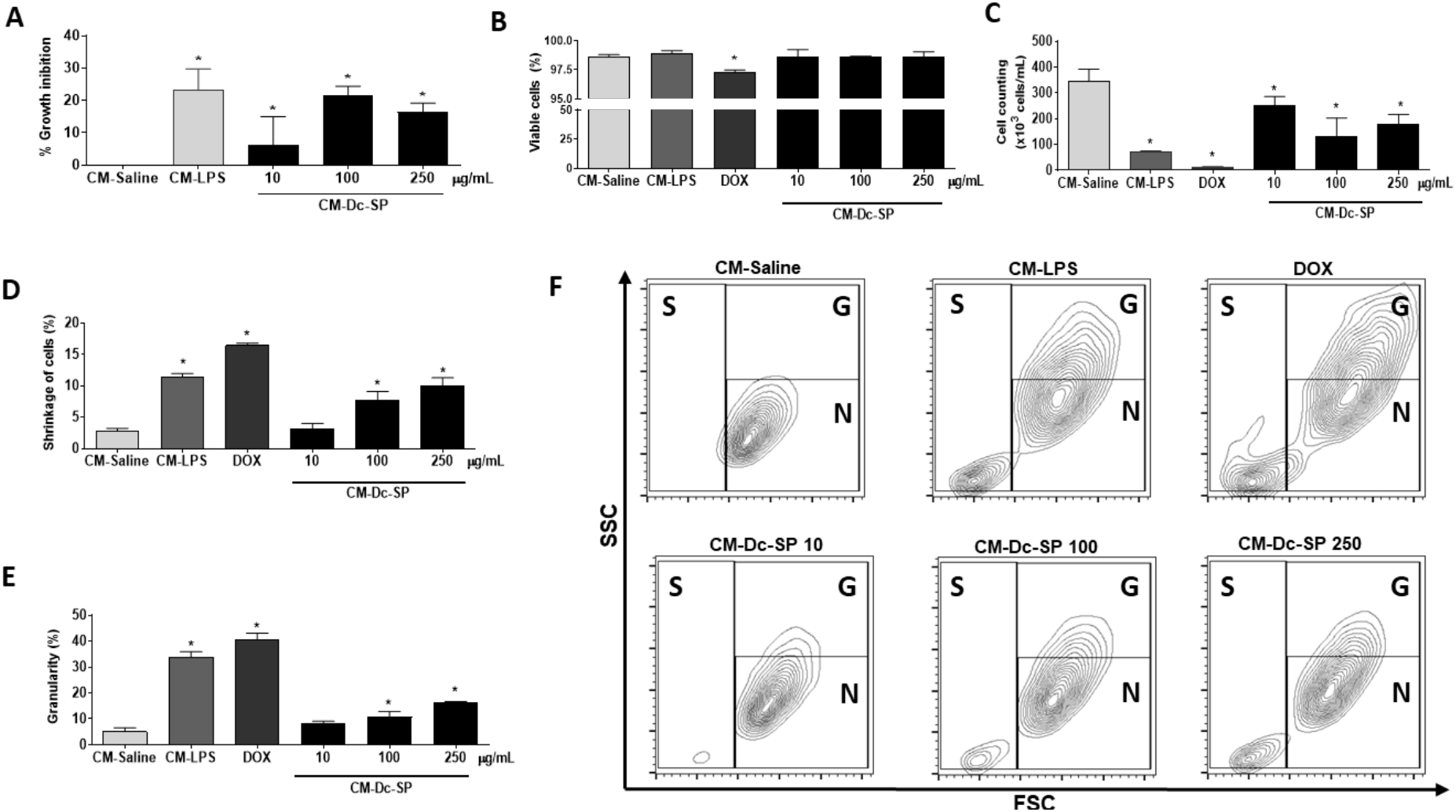
Antiproliferative effect of conditioned medium of macrophages stimulated with sulfated polysaccharide from *Dictyota caribaea* on melanoma. Conditioned medium (CM) was obtained from the supernatant of RAW264.7 cells incubated with saline (CM-Saline), *E. coli* lipopolysaccharide at 10 ng/mL (CM-LPS), doxorubicin at 0.6 µ M (DOX) and Dc-SP (CM-Dc-SP) for 24h. B16-F10 cells were then incubated with CM for 48h. **A**, Antiproliferative effect of Dc-SP against the murine metastatic melanoma cell line (B16-F10) evaluated by SRB assay. **B** and **C**, Cell count and cell viability of B16-F10 cells by flow cytometry. **D-F**, Cell morphology analyzed by flow cytometry. Data of A-E are presented as mean ±SD. Differences between data normalized to the percentage of cell growth inhibition of negative control versus other groups were determined by the analysis of variance (ANOVA) followed by Dunnett’s post-test. ^*^p<0.05. Legend: **S**, Cell shrinkage; **G**, Granulosity; **N**, Normal morphology of cells. Results are representative of two independent experiments, each one performed in triplicate.

A decrease in the number of cells can be observed at concentrations of 100 and 250 µg/mL of CM of Mφ activated by Dc-SP when compared with the control group (Fig. 3B), suggesting that Dc-SP favors an inhibition of B16-F10 murine melanoma proliferation. Doxorubicin (DOX) at 0.4 µM also reduced cell counts. However, as shown in Fig. 3C, no changes were found in the integrity of the membrane of cells treated with CM, showing that the sample did not damage the cell membrane. Only DOX changed the percentage of cells with intact membranes. Besides, the morphological analysis showed increases in the populations of cell shrinkage and cells with high granularity (Fig. 3D-F).

Engström and colleagues demonstrated that the CM of human Mφ polarized to an M1-phenotype after LPS and interferon-gamma (IFN-γ) exposure inhibits the proliferation of colorectal adenocarcinoma cell lines [43]. Already the CM from RAW264.7 polarized to M1-like phenotype with fucoidan from *L. japonica* significantly reduced the cell viabilities of two tumor cell lines (HCT116 and RKO) by about 40% [33]. In the same way, fucoidan from *Cladosiphon okamuranus* did not inhibit cell growth of sarcoma 180 *in vitro*. However, when cocultured with RAW264.7 in contact with these fucoidans, tumor cell growth was inhibited. This effect was attributed to the NO release by RAW264.7 as a marker for the M1 phenotype [44]. Our approach could not relate NO effect, once we used the supernatant, which already converted NO to nitrite and later is not active. Nevertheless, due to the recognized cytotoxic effect of the TNF family, we used tested the antiproliferative activity of TNF-α to evaluate its putative role on the B16-F10 inhibition by CM from RAW264.7 cells. However, TNF-α. did not inhibit B16-F10 cell growth after 48h of treatment at concentrations found in CM samples on ELISA assay. Therefore, other mediators produced by Mφ must be related to the inhibition of B16-F10 cell growth.

Apoptosis is one of the main mechanisms by which anticancer chemotherapeutic agents kill cells. Hydrophilic high molecular weight DNA markers, such as propidium iodide, cannot penetrate intact cells due to their size and do not label apoptotic cells unless they exhibit changes in plasma membrane permeability, as occurs in the late stages of apoptosis [45]. Cell shrinkage due to dehydration can be detected in the early stages of apoptosis as a decrease in the intensity of the forward scatter (FSC) light scattering signal [46,47]. Some fucans exhibit antiproliferative activity through various mechanisms, such as induction of cell cycle arrest, apoptosis, and activation of the immune system. These activities such as the induction of inflammation through the immune system, oxidative stress, and mobilization of stem cells have been reported to be linked to these anticancer properties [48].

One previous report suggested that the antitumor activity of fucoidan Dc-SP against S-180 cells *in vivo* is related to the enhancement of immune responses [13]. It shows that Dc-SP has an antitumor activity through immunomodulation of the tumor microenvironment rather than directly killing tumor cells, which could be promising in terms of reducing the side effects of cytotoxic substances.

Taken together, our data suggest that this antitumor activity *in vivo* previously reported for Dc-SP may be related to NO and cytokines production and M1 activation markers for M1 on Mφ as well.

## 4. Conclusions

Dc-SP polarizes Mφ to a pro-inflammatory and an antitumor phenotype *in vitro*.

## Acknowledgments

This work was supported by National Institute of Science and Technology on Biodiversity and Natural Products (INCT BioNat-CNPq/FAPESP, Finance code #465637/2014-0). The authors also would like to thank the Multi-User Facility of Drug Research and Development Center of Federal University of Ceará for technical support.

## Declaration of interest

The authors declare no conflict of interest.

## Author contributions

**Alexia Nathália Brígido Assef:** Methodology, Validation, Writing – Original Draft, Formal analysis, Investigation, Visualization; **Felipe Barros Teles:** Methodology, Writing – Review & Editing; **Leonardo Paes Cinelli**: Funding acquisition, Methodology, and Validation**; Diego Veras Wilke**: Conceptualization, Funding acquisition, Supervision, Validation, Writing – Review & Editing.

## Conflict of interest

None.

## Data Availability Statement

The data underlying this article will be shared on reasonable request to the corresponding author.

## Notes

### Competing Interest Statement

The authors have declared no competing interest.

